# Proline-2’-deoxymugineic acid (PDMA) increases seed quality and yield by alleviating iron deficiency symptoms in soybean under calcareous-alkaline conditions

**DOI:** 10.1101/2024.10.02.616232

**Authors:** Zahit Kaya, Amir Maqbool, Motofumi Suzuki, Emre Aksoy

## Abstract

Iron (Fe) deficiency in crops, particularly in calcareous-alkaline soils, poses a major challenge due to Fe immobilization. While synthetic chelators like EDTA and EDDHA are commonly used to improve Fe availability, proline-2’-deoxymugineic acid (PDMA) has emerged as a promising alternative, enhancing Fe nutrition in crops such as rice and cucumber. This study aimed to assess the effectiveness of PDMA on soybean growth and yield under calcareous-alkaline conditions. A pot trial demonstrated that PDMA improves Fe uptake, translocation, and storage in soybeans, leading to increased chlorophyll content, and enhanced root and shoot growth. Even at low dosages, PDMA significantly improved plant development, with the highest dosage (30 μM) resulting in notable increases in Fe, Zn, Mn, and Mg concentrations in roots, leaves, and seeds, surpassing the effects of the synthetic chelator Fe-EDDHA in several parameters. Additionally, 30 μM PDMA substantially boosted soybean yield, increasing pod and seed number, and 100-seed weight. It also improved seed quality by increasing protein and oil content. These findings suggest that PDMA offers a sustainable, effective alternative to traditional Fe chelators, providing a viable solution for addressing Fe deficiency and enhancing crop biofortification in challenging soil conditions.

## Introduction

Iron (Fe) ranks as the third most restricting factor for plant development, mainly due to its limited solubility in aerobic conditions and calcareous soils, despite its abundance in the soil (Kobayashi 2019). In fact, over one-third of soils globally are classified as Fe-deficient because of high soil pH levels. This deficiency impacts both prokaryotic and eukaryotic organisms, which require Fe for essential functions, including the activity of (leg)haemoglobin and enzymes involved in DNA and RNA synthesis, nitrogen fixation, and immune responses (Li 2023). A reduction in these critical activities leads to Fe deficiency chlorosis (IDC), resulting in stunted growth and significant yield losses. Consequently, crops accumulate lower levels of Fe in their edible parts when grown under Fe-deficient conditions. The consumption of such crops, with lower Fe content in their seeds, directly affects human health. Fe deficiency is a major global health issue, often referred to as “hidden hunger,” impacting approximately 2 billion people worldwide (Szerement et al., 2022). It is the primary cause of Fe deficiency anemia, a condition with high prevalence, particularly in low- and middle-income countries (WHO, 2020). Uneven distribution of nutrients, often with higher Fe content in non-edible parts of crops, highlights the need for effective strategies to combat micronutrient malnutrition. Among various strategies to mitigate Fe deficiency, agronomic biofortification stands out due to its easy application, cost-effectiveness, and minimal adverse effects on crop yield (Szerement et al., 2022). Therefore, it is a sustainable strategy for increasing crop yields and nutritional quality in short term. Recent advances in the development of new fertilizer forms and emerging techniques have further enhanced the efficiency of agronomic biofortification. These innovations aim to improve the uptake of micronutrients by crop plants, thereby addressing challenges related to human nutrition and health.

To cope with Fe deficiency, plants have developed two main mechanisms, namely “Strategy I” and “Strategy II”, to obtain this essential nutrient from the soil (Ning et al., 2023). In Strategy I, primarily observed in dicots such as soybean and non-grass monocots, ferric iron (Fe^+3^) is reduced to ferrous iron (Fe^+2^) by ferric chelate reductase (FCR) localized on the plasma membrane of root epidermal cells, thereby enhancing Fe bioavailability (Li et al., 2023). Then, Fe^+2^ is carried into plant roots by specific transporters, including natural resistance-associated macrophage proteins (NRAMPs) and Fe-regulated transporters (IRTs). This process is regulated by FER-like Fe deficiency-induced transcription factor (FIT) (Rajniak et al., 2018). On the other hand, in Strategy II plants, such as *Poaceae*, a group of compounds known as mugineic acid family phytosiderophores (MAs) are released into the rhizosphere (Li et al., 2023). These MAs serve to bind and chelate Fe^3+^ in the soil. The resulting MAs-Fe^3+^ complex is then absorbed by the plant roots through transporters referred to as yellow stripe (YS) transporters (Ning et al., 2023).

Strategy I plants respond to Fe deficiency primarily in high-pH conditions and are often found in calcareous soil (Zuo and Zhang, 2011). Recent research has shown that these plants can enhance their Fe nutrition by utilizing a compound called 2’-deoxymugineic acid (DMA), which is produced by Strategy II plants (Suzuki et al., 2016). Strategy I plants also possess transporters similar to those involved in DMA uptake (Ding et al., 2009; Xiong et al., 2013). However, when it comes to practical agricultural use, employing DMA as an Fe chelator to improve the Fe nutrition of Strategy I plants is not feasible due to its tendency to degrade easily in soil, and the high cost associated with its production (Suzuki et al., 2021). Recently, a new synthetic phytosiderophore called proline-2′-deoxymugineic acid (PDMA) has emerged as a promising Fe chelator to increase Fe uptake by plants (Suzuki et al., 2021). PDMA differs from natural DMA in its structure, as it replaces L-azetidine with L-proline, making it more stable to biodegradation by soil microorganisms and cost-effective to produce. When applied to rice plants in calcareous soil, Fe-PDMA complex enhanced Fe uptake more efficiently than chelated forms of Fe, namely Fe-EDDHA and Fe-EDTA (Suzuki et al., 2021). The improved availability of PDMA can be attributed to its direct absorption by YSL transporters in rice, maize and barley. Different stereoisomers of PDMA were shown to be transported by YSL transporter in a yeast complementation study while only some specific forms could rescue IDC symptoms in rice in calcareous soil (Namba et al., 2024). The application of metal-free PDMA to calcareous soil also led to recovery from Fe deficiency. Supplementation of Fe-PDMA alleviated the IDC symptoms more effectively than Fe-EDDHA by increasing Fe concentrations in the aboveground tissues of both *Poaceae* but also dicots, such as cucumber, pumpkin (Ueno et al., 2021), and peanut (Wang et al., 2023). These findings consistent with altered FCR activity in plant roots under alkaline conditions, indicating a strong potential for PDMA to enhance both yield and seed Fe content in dicots (Ueno et al., 2019). However, its impacts on seed quality have not been studied in plants before.

Soybean (*Glycine max* (L.) Merrill), being a major global crop, holds significant potential for biofortification efforts. In 2022, 348.8 million tons of soybeans were produced worldwide, with the United States contributing significantly to this total. The U.S. production was valued at $59.2 billion and accounted for almost 60% of the world’s oilseed production (FAOSTAT, 2023). Although it is the most important oil seed crop grown extensively across the world, soybean yield and Fe content can be adversely affected by high pH soils, which promote IDC (Merry et al., 2022). IDC is a significant issue in calcareous soils, including the northern Midwest United States, impacting approximately 24% of the soybean crop and leading to considerable yield losses (Merry et al., 2022). Soybean was proposed as a model organism to study the effects of Fe deficiency and fertilization (Aksoy et al., 2017). In modern agriculture, synthetic Fe-chelating compounds like EDTA and EDDHA are commonly utilized as stabilizers in Fe fertilizers to address Fe deficiency in crops. Due to the alkaline pH of calcareous soils, Fe fertilizers tend to become immobilized in the soil after application (Abadía et al., 2011). Nevertheless, PDMA has shown a stronger capability to enhance Fe nutrition in rice (Suzuki et al., 2021) and cucumber (Ueno et al., 2021). However, more research is needed to understand the impact of PDMA on seed quality and yield in soybean cultivation in calcareous soil. Therefore, in the present study, we aimed to assess the effectiveness of PDMA when applied to soybeans in a pot trial under calcareous-alkaline conditions. We found that it effectively enhances Fe uptake, translocation and storage, leading to increased soybean yields, seed quality and micronutrient concentrations. These findings indicate promising prospects for the use of PDMA in agricultural systems and PDMA could serve as a viable alternative to conventional synthetic chelates to alleviate the Fe deficiency symptoms and yield losses in soybeans while increasing the seed Fe content for utilization in biofortification.

## Materials and Methods

### Plant materials and growth conditions

A local soybean cultivar (*Glycine max* L.), Atakişi, was used in this study due to its high sensitivity to Fe deficiency (Maqbool, 2018). After surface sterilization by washing the seeds in 0.25 % sodium hypochlorite for 5 mins, followed by 3 times washing with distilled water, the seeds were germinated in vitro on Petri plates for five days in between wet tissue papers in a growth chamber at 25 ± 1 °C and 60% relative humidity. After five days, germinated seedlings with similar root lengths (at similar growth stages) were transferred to the potting mixture (soil, peat and perlite mixture at a ratio of 2:2:1). The soil was taken from Niğde/Türkiye and its properties are provided in Supplemental Table 1. Plants were grown in a climate-controlled greenhouse in pots of 28 cm diameter at 25 ± 1 °C (day) and 20 ± 1 °C (night) and 50 % humidity under long-day conditions (16:8 h of day:night) supplemented with 580 ± 75 μmol PAR m^-2^ s^-1^. Each pot was filled with 5 kg of potting mixture and the experiment was set up according to a completely randomized design, where each pot represented one biological replicate, and a total of twelve replications were used for each treatment. Four plants were grown in each pot to represent technical repeats.

In one set of the pots (normal pH), the pH of the potting mixture was measured as 5.7 according to Baker et al. (1983). This treatment was defined as Control + (C +) and did not receive any other applications. In the second set, Fe deficiency was applied by increasing the pH of the potting mixture to 8.7 by mixing thoroughly 9 g of calcium oxide (CaO, Sigma) / kg soil. This treatment was defined as Control - (C -). Then, different dosages of proline-2’-deoxymugineic acid (PDMA) (AICHI Steel Corporation, Japan) (3 µM, 15 µM, and 30 µM in double-distilled water) were applied to the plants grown in calcareous-alkaline soil with pH of 8.7. 30 µM of 6% Fe-EDDHA (4.8% o-o) (Syngenta) was used as a fertilizer control (Schenkeveld et al., 2008). When plants reached the vegetative 1 (V1) growth stage (around 15 days after emergence), 150 mL of PDMA or Fe-EDDHA were applied to the rhizosphere of each plant 4 times in 5-day intervals. Plants in C + and C - sets received the same volume of double-distilled water. All physiological, biochemical, and molecular measurements were taken from six random pots when the plants reached V3 growth stage (on day 35 after emergence). Remaining pots received regular irrigation (around 600 mL of double-distilled water in every three days) until maturity. The water content of the potting mix was measured in every two days to ensure 85% of field capacity was assured (Signorelli et al., 2022).

### Physiological measurements

The chlorophyll index (Soil Plant Analysis Development – SPAD - value) was measured from the first fully developed trifoliate leaves by a SPADmeter (Minolta). Each data was collected from at least three independent measurements from each plant. Then, the first trifoliate leaves were photographed using a digital camera to asses the leaf area using ImageJ software (http://rsbweb.nih.gov/ij/) (Aksoy et al., 2012). The root and shoot lengths were measured by a tape. Following imaging, the fresh weights of the roots, shoots, and leaves were determined by using a digital balance. Subsequently, these plant parts were dried in an oven at 65 °C for 24 hours, and their respective dry weights were recorded.

To determine the chlorophyll content, the fresh weight (FW) of the first trifoliates were recorded. Subsequently, total chlorophyll was extracted using a mortar and pestle in 10 mL of 80% (v/v) acetone and the resulting mixtures were kept overnight at 4 °C. The next day, the samples were centrifuged at 13,000 g at 4 °C for 5 minutes and the absorbance of the supernatant at 470, 646.8, and 663.2 nm was read against 80% acetone as a blank. The total chlorophyll content (chla + chlb) was calculated using the formula (7.15 x A663.2 + 18.71 x A646.8) / 1000 / fresh leaf weight (Lichtenthaler, 1987).

### Ferric chelate reductase (FCR) activity

The FCR activity in the roots was assessed according to Aksoy and Koiwa (2013). Briefly, approximately 100 mg of the sample was collected from the area close to the root tip. The roots were washed three times with distilled water after carefully separated from the potting mix and their fresh weights were recorded. Then, root samples were incubated in the FCR enzyme activity solution (0.1 mM Fe^3+^-EDTA (Sigma) and 0.3 mM Ferrozine (Sigma)) for 12 hours in the dark at room temperature and the absorbance of the solution was read at 562 nm against a blank lacking root. Enzyme activity was calculated using a molar extinction coefficient of 28.6 mM^-1^ cm^-1^ and normalized based on the root fresh weight.

### Rhizosphere acidification

Rhizosphere acidification was assessed according to (Pizzio et al., 2015). Briefly, the roots were washed three times with distilled water after carefully separated from the potting mix and their fresh weights were recorded. Then, the roots were soaked in 30 mL of ½ Hoagland’s solution (Hoagland and Arnon, 1938) supplemented with 2 mM MES buffer (pH 6.8) and incubated for 24 hours in a growth room at 25 ± 1 °C (day) and 20 ± 1 °C (night) and 50 % humidity under long-day conditions (16:8 h of day:night) supplemented with 580 ± 75 μmol PAR m^-2^ s^-1^. An assay solution without plants was also incubated to serve as a control. Crude H^+^ released to the medium was calculated from pH=-log [H^+^] and normalized based on the root fresh weight.

### Metal concentration analysis in the leaves, roots, and seeds

Roots, first trifoliate leaves, and seeds were collected and pooled together from three biological replicates for each organ. Samples were incubated in 2 mM CaSO_4_ (Sigma) and 10 mM EDTA (Sigma) for 10 minutes followed by washing thrice with distilled water to eliminate any metal particles attached to the sample surface. The samples were divided into 3 technical repeats of 100 mg and dried in acid-resistant borosilicate test tubes for 48 hours at 65°C (Aksoy et al., 2012). Then, the samples were digested in 3.25 mL of 15 M nitric acid (HNO_3_) (Sigma), and 2 mL of 30% hydrogen peroxide (H_2_O_2_) (Sigma) at 100 °C for 1 hour, 150 °C for 1 hour, 180 °C for 1.5 hours, and lastly at 210 °C until no liquid is left in the test tubes (Vasconcelos et al., 2006). Finally, the samples were re-dissolved with 3.25 mL of 15 M HNO_3_, and 2 mL of 30% H_2_O_2_ and the Fe, Zn, Mn, and Mg levels were determined in Inductively Coupled Plasma Mass Spectrometry (ICP-MS) (Bruker Aurora M90). Each sample was read five times in the pulse detector mode.

### Yield parameters

The yield measurements were taken at the crop maturity stage after harvesting of the seeds. Pod and seed number per plant and 100-seed weight was recorded (Çalişkan et al., 2007).

### Seed protein and oil content

Seed total protein content was determined by the Kjeldahl apparatus (Çalı kan et al., 2018). Oil content was measured from well-ground seeds according to the method described by Am, A.A.P., 2005 by using the ANKOM extractor (Çalı kan et al., 2018).

### Gene expression analyses

The total RNA was extracted from the roots using TRIzol reagent (Sigma) (Chomczynski and Sacchi, 1987). Following the removal of genomic DNA by DNase I treatment, 1 μg total RNA was converted to first strand cDNA using SuperScript® VILO ™ cDNA synthesis kit (Invitrogen). Reverse transcribed products were analyzed by Real-Time qPCR (Qiagene) in a reaction of 20 ng of cDNA mixed with 12.5 µL of RealQ Plus 2x Master Mix (Ampliqon) and 0.8 µM of the specific primers in a final volume of 25 µL. The expressions of *GmIRT1, GmFRO2, GmFRD3*, and *GmFIT* was determined (Santos et al., 2015) (Table 1). *GmELONGATION FACTOR 1-BETA* (*GmEFL1B*) was used as an internal control to normalize the data (Jian et al., 2008). Each reaction was run in two technical replicates and data is analyzed according to the 2^-ΔΔCt^ method as described by (Rao et al., 2013).

**Table 1.** Zn, Mn, and Mg concentrations in the roots, and trifoliate leaves. Data are represented as means ± SEM (n = 3). Different letters indicate a significant difference (p < 0.05) using ANOVA and Tukey’s Tests.

| Treatments | Zn ($\mu\text{g/g DW}$) | Mn ($\mu\text{g/g DW}$) | Mg ($\mu\text{g/g DW}$) |
| --- | --- | --- | --- |
| **Root** |  |  |  |
| C + | 23.5 ± 0.6 b | 15.5 ± 0.2 c | 380 ± 4.6 c |
| C - | 17.4 ± 0.8 c | 13.8 ± 0.5 d | 298 ± 8.8 e |
| PDMA 3 | 18.0 ± 0.8 c | 15.0 ± 0.2 cd | 300 ± 4.3 e |
| PDMA 15 | 17.0 ± 0.7 c | 19.0 ± 0.2 b | 330 ± 2.0 d |
| PDMA 30 | 29.5 ± 0.8 a | 19.5 ± 0.2 b | 505 ± 1.9 a |
| Fe-EDDHA 30 | 21.5 ± 0.2 b | 21.5 ± 0.2 a | 405 ± 2.2 b |
| **Trifoliolate** |  |  |  |
| C + | 4.6 ± 0.3 b | 11.5 ± 0.2 b | 235 ± 3.9 b |
| C - | 3.3 ± 0.4 bc | 8.3 ± 0.4 d | 202 ± 2.8 c |
| PDMA 3 | 2.8 ± 0.2 c | 9.6 ± 0.2 cd | 205 ± 2.1 c |
| PDMA 15 | 3.1 ± 0.4 bc | 9.8 ± 0.6 bcd | 210 ± 2.1 c |
| PDMA 30 | 14.0 ± 0.4 a | 10.5 ± 0.2 bc | 270 ± 4.5 a |
| Fe-EDDHA 30 | 3.8 ± 0.3 bc | 16.0 ± 0.2 a | 215 ± 2.0 c |

### Statistical analysis

All data were analyzed with the R software (R Core Team, 2021) via one-way ANOVA followed by Tukey’s post-hoc test to indicate and determine the significant differences between treatment groups. Illustrations and graphs were drawn in Microsoft Excel.

## Results

### Changes in physiological parameters with PDMA application under alkaline conditions

After 20 days of fertilizer application, distinct differences in chlorosis were observed among the treatment groups. Notably, chlorosis was evident in the trifoliates of the plants grown in calcareous-alkaline conditions, where pH was 8.5 (C -) and the ones treated with 3 μM PDMA whereas the remaining PDMA treatments and Fe-EDDHA showed no signs of chlorosis (Fig. 1a). Analysis of chlorophyll index further supported these observations, with 15 μM PDMA treatment demonstrating the highest SPAD value. Conversely, Control (-) and 3 μM PDMA treatments exhibited the lowest SPAD values (Fig. 1b). 15 μM PDMA application could increase the chlorophyll index significantly by 38% compared to the plants grown in calcareous-alkaline potting mix. Chlorophyll a and chlorophyll b contents of the plants grown in calcareous-alkaline conditions and the ones treated with 3 or 15 μM PDMA were significantly lower than the Control (+) plants while treatment with 30 μM PDMA or Fe-EDDHA returned the chlorophyll a and b contents back to Control (+) levels (Fig. 1c and 1d). These results highlight a correlation between treatments and chlorophyll parameters, with high PDMA dosages consistently showing high chlorophyll values and Control (-) and 3 μM PDMA the lowest. Overall, these observations suggest a dose-dependent effect of PDMA application on the alleviation of Fe-deficiency chlorosis.

**Figure 1.**
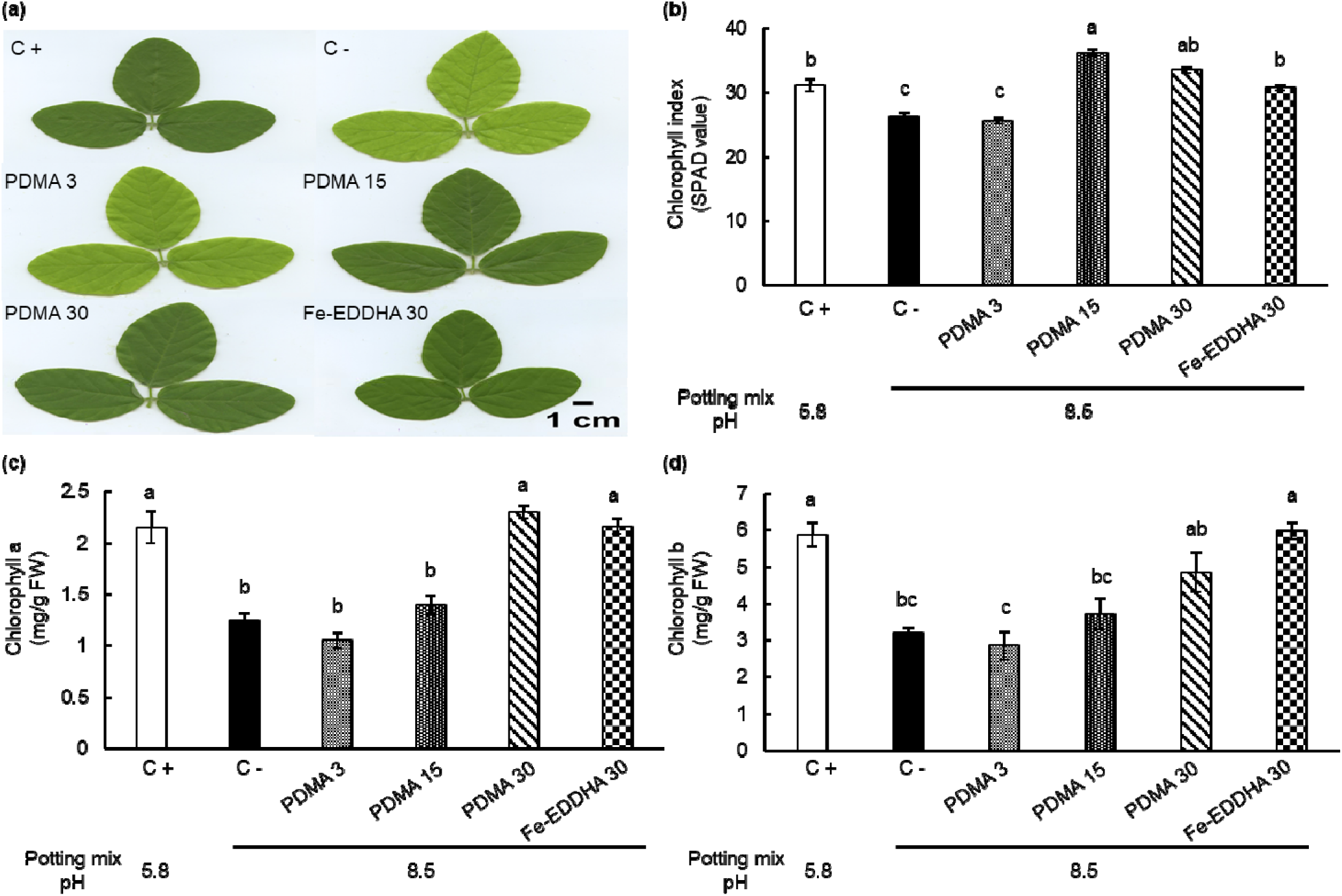
Effects of PDMA application on leaf chlorosis of soybean grown in calcareous-alkaline conditions. (a) Trifoliate images, (b) Chlorophyll index, (c) Chlorophyll a content, (d) Chlorophyll b content. pH conditions of the potting mix are indicated for each application in the bar graphs. Data are represented as means ± SEM (n = 6). Different letters indicate a significant difference between treatments (p < 0.05) using ANOVA and Tukey’s post-hoc test.

To assess the effects of PDMA application on plant development, we measured the root and shoot lengths, root and trifoliate fresh and dry weights as well as trifoliate surface area. Plant roots significantly decreased by 35% (p < 0.05) when grown in calcareous-alkaline potting mix compared to the Control (+) while they could not be rescued significantly by 3 μM PDMA or 30 μM Fe-EDDHA applications (Fig. 2a). However, 30 μM PDMA application to the plants grown in calcareous-alkaline potting mix increased the root length significantly by 50% (p < 0.05) and reached to the levels observed in Control (+) plants. A similar trend was observed in the shoot lengths (Fig. 2b). Plant shoots were 36% shorter when grown in calcareous-alkaline conditions compared to the Control (+) whereas 30 μM PDMA application substantially increased the shoot lengths back to the Control (+) levels. Although 30 μM Fe-EDDHA application could improve the shoot length by 14% in the plants grown in calcareous-alkaline conditions, it was not able to make a significant improvement.

**Figure 2.**
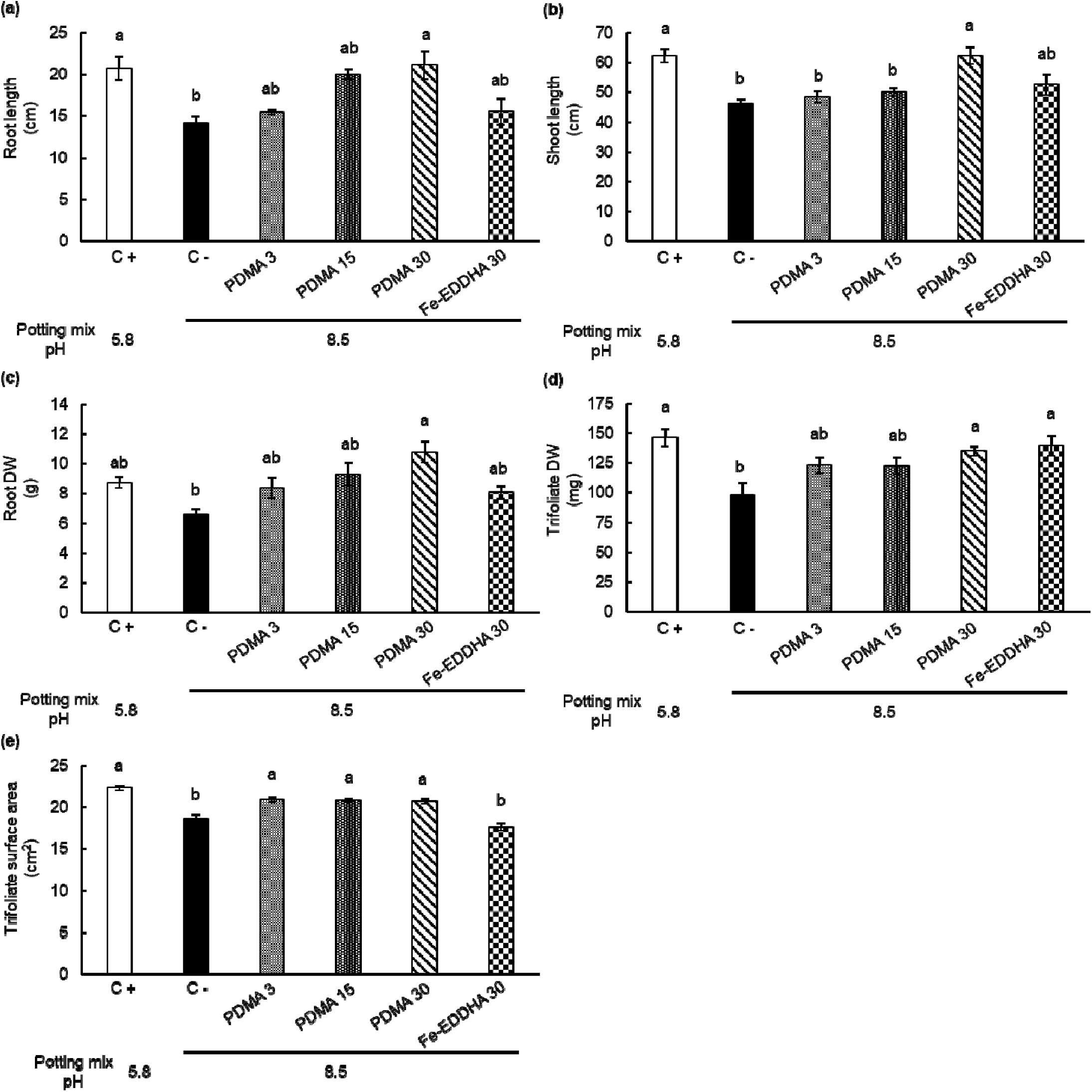
Effects of PDMA application on growth parameters of soybean grown in calcareous-alkaline conditions. (a) Root length, (b) Shoot length, (c) Root dry weight, (d) Trifoliate dry weights, (e) Trifoliate surface area. pH conditions of the potting mix are indicated for each application in the bar graphs. Data are represented as means ± SEM (n = 6). Different letters indicate a significant difference (p < 0.05) using ANOVA and Tukey’s Tests.

Root dry weight results were consistent with the root lengths, where increasing PDMA dosage increased the root dry weight when the plants were grown in calcareous-alkaline conditions reaching to the maximum in 30 μM of PDMA application (Fig. 2c). Increasing PDMA dosage applications increased the trifoliate dry weight and surface area compared to the Control (-) group, returning back to the Control (+) levels (Fig. 2d and Fig. 2e). Moreover, 30 μM Fe-EDDHA application increased the trifoliate dry weight, rescuing it back to the Control (+) levels although it did not have any positive effects on the trifoliate surface area. These findings emphasize the complex effects of PDMA and Fe supplementation on soybean growth parameters, highlighting the intricate interplay between nutrient availability and plant physiological responses in calcareous soil conditions.

### Effect of PDMA on FCR activity

Ferric chelate reductase (FCR) serves as the major enzyme governing Fe uptake in dicots (Satbhai et al., 2017). The measurement of its activity in plant roots is essential for comprehending the potential impact of PDMA application. FCR activity was significantly increased by 71% (p < 0.05) when grown in calcareous-alkaline potting mix compared to the Control (+) (Fig. 3a). Application of 3 μM, 15 μM and 30 μM PDMA in calcareous-alkaline conditions significantly decreased the FCR activity by 22%, 26% and 48%, respectively (p < 0.05). The FCR activity levels returned back to the Control (+) conditions only after 30 μM PDMA application while 30 μM Fe-EDDHA application could only decrease the FCR activity by 30%. These results suggest that the application of higher dosages of PDMA and Fe-EDDHA effectively mitigates the negative effects of alkalinity-based Fe deficiency on FCR activity, bringing it back to levels observed under control conditions.

**Figure 3.**
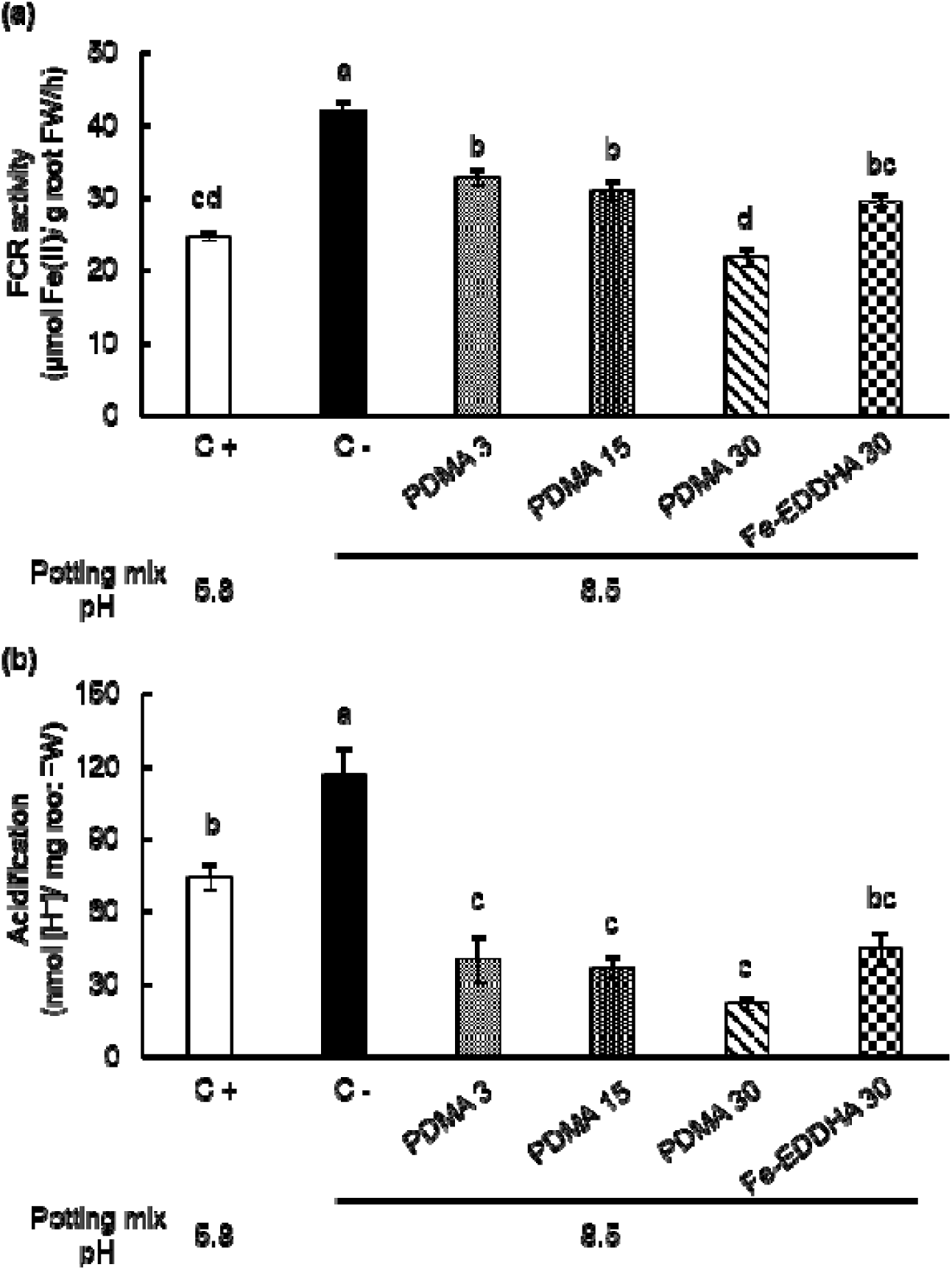
Effect of PDMA application on FCR activity and rhizosphere acidification of soybean grown in calcareous-alkaline conditions. pH conditions of the potting mix are indicated for each application in the bar graphs. Data are represented as means ± SEM (n = 6). Different letters indicate a significant difference (p < 0.05) using ANOVA and Tukey’s Tests.

Plants can release protons into the soil to increase the Fe^3+^ solubility and availability by decreasing the pH (Vélez-Bermúdez and Schmidt, 2023). Plants acidified the rhizosphere significantly higher by 57% (p < 0.05) under Control (-) conditions compared to the Control (+) (Fig. 3b). However, application of Fe-EDDHA effectively decreased the acidification levels back to the Control (+) levels while application of PDMA caused significant decreases below the Control (+) levels even under low dosages. Interestingly, the pH of the potting mix did not change after PDMA applications, suggesting that PDMA application can alter the ability of plants to acidify the rhizosphere.

### Effects of PDMA on the expression of known Fe uptake genes

We also examined the expression of known Fe uptake genes in the roots of soybean plants treated with PDMA while grown in calcareous-alkaline conditions (Fig. 4). We observed that *GmFIT, GmIRT1, GmFRO2*, and *GmFRD3* were all significantly induced in the roots under high soil pH conditions. However, as the dosage of PDMA increased, the expression levels of these genes gradually decreased and eventually reached the levels observed in the Control (+) treatment at the highest PDMA application. Although Fe-EDDHA application also restored the expression levels, it did not reduce the expression levels of *GmFIT* and *GmIRT1* to the levels of the Control (+) treatment. Indeed, the expression levels of *GmFIT* and *GmIRT1* were reduced by 53% and 48%, respectively after Fe-EDDHA application while this decreased increased to 76% after 30 μM PDMA application compared to the Control (-) (p < 0.05). On the contrary, both 30 μM PDMA and Fe-EDDHA applications significantly decreased the expression levels of *GmFRO2*, and *GmFRD3* back to the Control (+) conditions. Overall, these findings suggest that PDMA application is selectively more effective in addressing Fe deficiency responses than does Fe-EDDHA at the molecular level in soybean roots.

**Figure 4.**
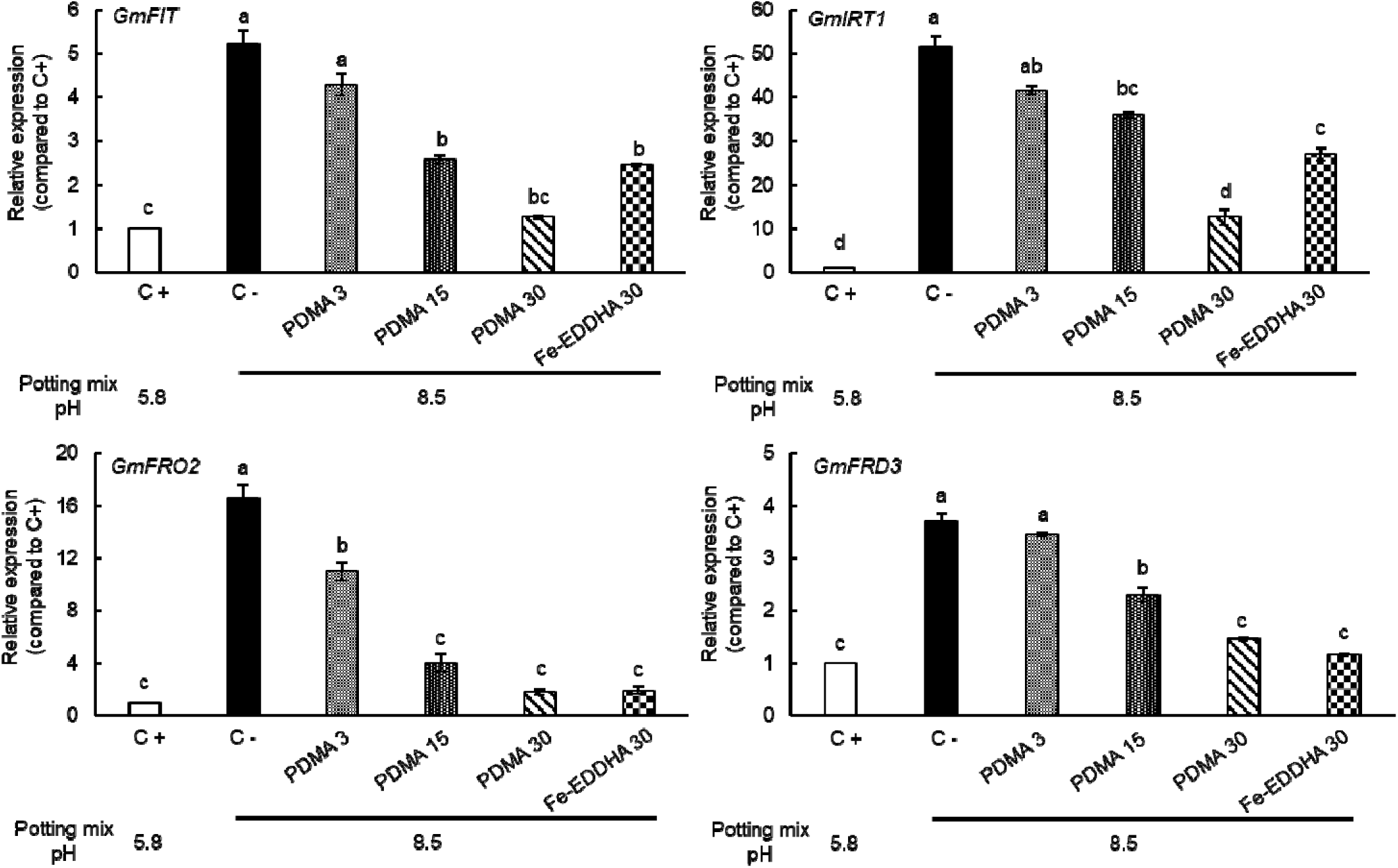
Effects of PDMA application on the expression levels of *GmFIT, GmIRT1, GmFRO2,* and *GmFRD3* in soybean grown in calcareous-alkaline conditions. pH conditions of the potting mix are indicated for each application in the bar graphs. Data are represented as means ± SEM (n = 3). Different letter(s) indicate a significant difference (p < 0.05) using ANOVA and Tukey’s Tests.

### Effects of PDMA on micronutrient levels in the roots, and trifoliate leaves

To assess the impact of different treatments on micronutrient uptake and accumulation in soybean plants following PDMA application, the concentrations of Fe, Mn, Zn, and Mg were examined in both the roots and leaves. Results revealed diverse effects of PDMA applications on Fe accumulation in different plant organs. In plants grown under calcareous-alkaline conditions, root Fe concentrations significantly decreased by 36% (p < 0.05) compared to the Control (+) condition (Fig. 5a). 3 μM PDMA application resulted in root Fe concentrations similar to those observed under the Control (-) treatment, while higher PDMA dosages led to significant increases in root Fe concentration. However, even at the highest PDMA dosage, root Fe levels did not reach those observed under Control (+) conditions. In contrast, treatment with Fe-EDDHA resulted in the highest root Fe concentrations, with levels that were 89% higher than those observed under Control (-) conditions and even 20% higher than under Control (+) conditions.

**Figure 5.**
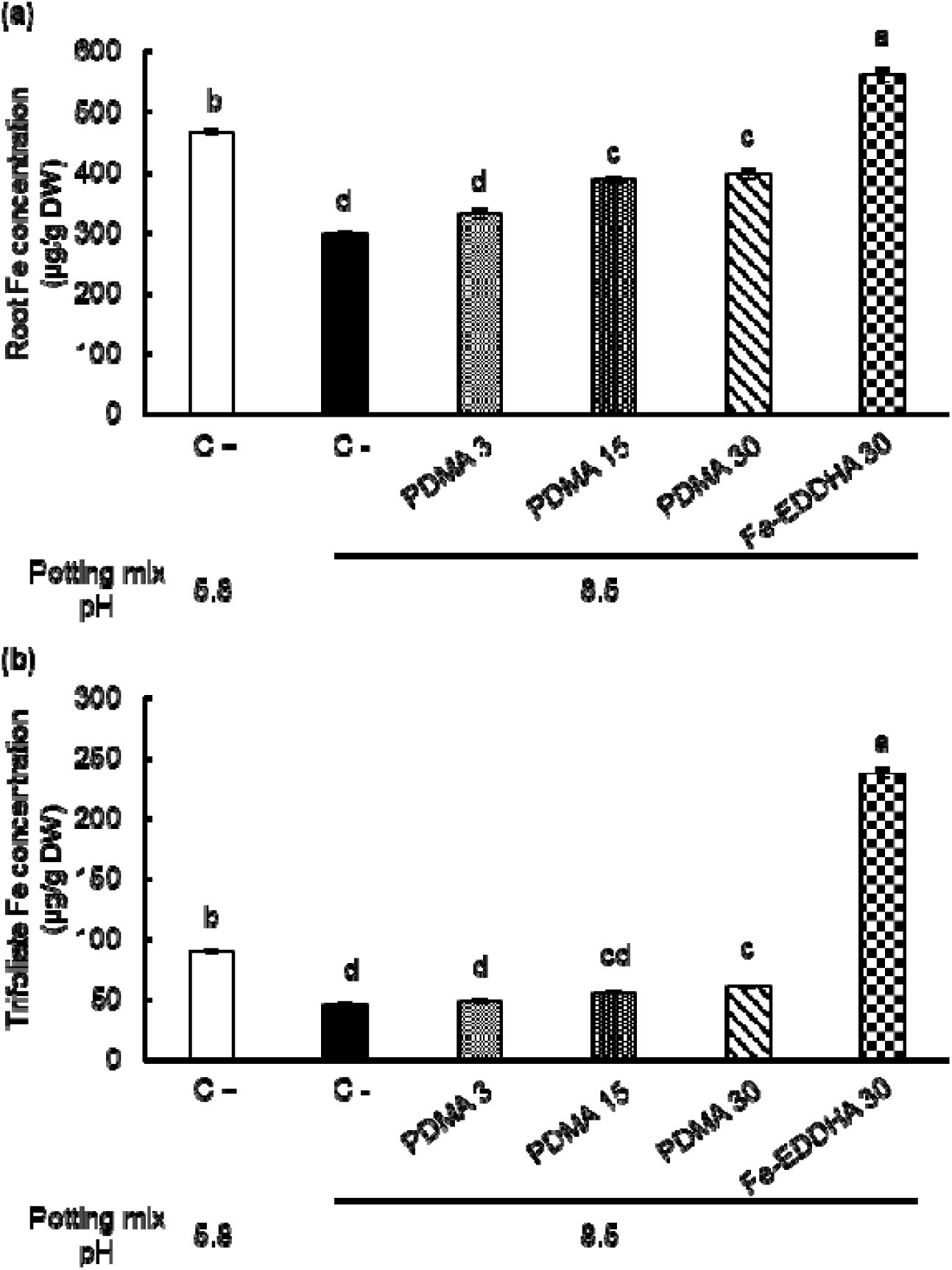
Effects of PDMA application on the Fe concentrations in the roots and trifoliate leaves of soybean grown in calcareous-alkaline conditions. (a) Root, (b) Trifoliate leaves. pH conditions of the potting mix are indicated for each application in the bar graphs. Data are represented as means ± SEM (n = 3). Different letters indicate a significant difference (p < 0.05) using ANOVA and Tukey’s Tests.

Analysis of leaf Fe concentrations also revealed significant differences among the treatments. The application of 30 μM Fe-EDDHA resulted in a notably higher leaf Fe concentration compared to all other groups, showing a 5.2-fold increase compared to Control (-) and a 2.6-fold increase compared to Control (+) conditions (Fig. 5b). Conversely, PDMA applications at 3 and 15 μM resulted in slight, nonsignificant increases in leaf Fe concentration, while the 30 μM PDMA application led to a significant 33% increase compared to the Control (-) condition. However, even at the highest PDMA dosage, leaf Fe concentration did not recover to Control (+) levels. Taken together, these findings indicate that Fe-EDDHA is a more effective Fe chelator than PDMA for enhancing root and leaf Fe concentrations, particularly in plants grown under calcareous-alkaline conditions. Fe-EDDHA not only restored Fe levels but also elevated them beyond those observed under Control (+) conditions.

Distinct patterns were observed in the concentrations of other divalent metals in plant organs following PDMA application. Root Zn concentrations decreased significantly by 26% (p < 0.05) when plants were grown under calcareous-alkaline conditions compared to the Control (+) condition (Table 1). 3 and 15 μM PDMA applications did not restore root Zn levels to those observed under Control (+) conditions. However, treatment with 30 μM Fe-EDDHA successfully restored root Zn levels to Control (+) conditions. In contrast, 30 μM PDMA application increased root Zn levels beyond those of Control (+), resulting in a significant 25% and 70% (p < 0.05) higher Zn accumulation in the roots compared to Control (+) and Control (-) conditions, respectively. A similar pattern was observed in trifoliate Zn concentrations. The application of 30 μM PDMA increased trifoliate Zn concentrations beyond Control (+) levels, while 30 μM Fe-EDDHA treatment did not cause a significant change in trifoliate Zn concentrations.

Mn concentrations decreased significantly by 11% and 28% (p < 0.05) in plant roots and trifoliates, respectively, when grown under calcareous-alkaline conditions compared to the Control (+) condition (Table 1). 3 μM PDMA application was sufficient to restore root Mn concentrations to Control (+) levels, while higher dosages significantly increased Mn levels beyond Control (+). Treatment with 30 μM Fe-EDDHA resulted in the highest root Mn concentrations, which were 38% and 56% higher (p < 0.05) than those observed under Control (+) and Control (-) conditions, respectively. In contrast to the effect of PDMA on root Mn levels, even the highest PDMA dosage barely restored trifoliate Mn concentrations to Control (+) levels. However, similar to the root results, the 30 μM Fe-EDDHA treatment significantly increased trifoliate Mn concentrations by 39% and 93% (p < 0.05) compared to Control (+) and Control (-) conditions, respectively.

Root and trifoliate Mg concentrations followed a similar trend, where higher PDMA dosages significantly increased Mg levels beyond Control (+) conditions. Among the treatments, the 30 μM PDMA application resulted in the highest Mg concentrations in both roots and trifoliates, while the 30 μM Fe-EDDHA treatment failed to restore trifoliate Mg concentrations to Control (+) levels. Taken together, these results suggest that higher PDMA dosages enhance divalent metal concentrations in plants grown under calcareous-alkaline conditions, particularly in the roots.

## Effects of PDMA on soybean yield parameters

Next, the impact of PDMA on soybean yield parameters was assessed across different treatment concentrations. Although yield parameters decreased when plants were grown under calcareous-alkaline conditions, there were no significant differences between the Control (+) and Control (-) conditions (Table 2). However, even the lowest dosage of PDMA application restored the yield parameters to levels comparable to the Control (+) condition. Similarly, 30 μM of Fe-EDDHA treatment successfully recovered the yield parameters to Control (+) levels. Furthermore, the application of 30 μM PDMA resulted in the highest yield parameters, significantly increasing the pod number per plant, seed number per plant, and 100-seed weight by 140%, 86%, and 54% (p < 0.05), respectively, compared to the Control (-) condition. Overall, these findings indicate that lower dosages of PDMA application are essential for increasing yield under calcareous-alkaline conditions, similar to Fe-EDDHA treatment, while higher PDMA dosages can lead to even greater yield improvements.

**Table 2.**
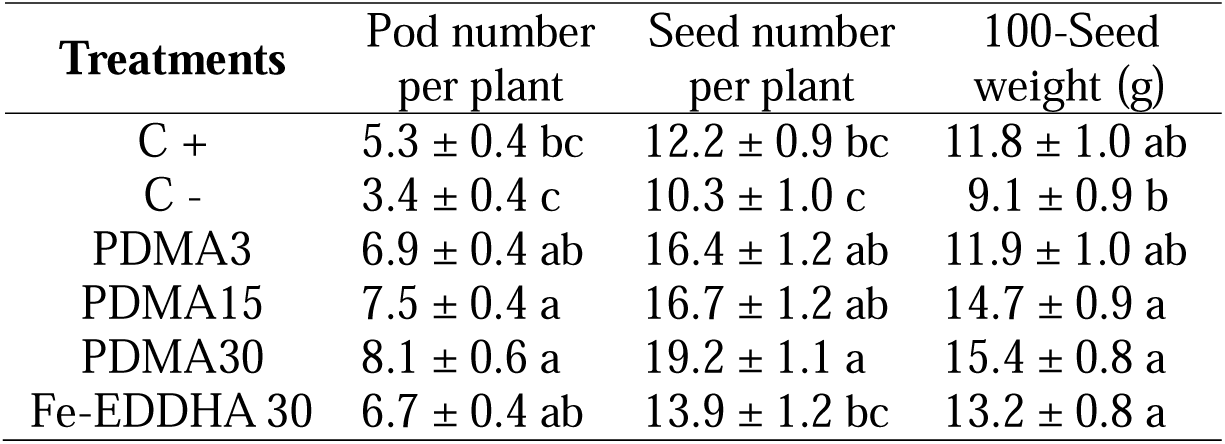
Effects of PDMA on soybean yield parameters. Data are represented as means ± SEM (n = 6). Different letters indicate a significant difference (p < 0.05) using ANOVA and Tukey’s Tests.

| **Treatments** | Pod number<br>per plant | Seed number<br>per plant | 100-Seed<br>weight (g) |
| --- | --- | --- | --- |
| C + | $5.3 \pm 0.4$ bc | $12.2 \pm 0.9$ bc | $11.8 \pm 1.0$ ab |
| C - | $3.4 \pm 0.4$ c | $10.3 \pm 1.0$ c | $9.1 \pm 0.9$ b |
| PDMA3 | $6.9 \pm 0.4$ ab | $16.4 \pm 1.2$ ab | $11.9 \pm 1.0$ ab |
| PDMA15 | $7.5 \pm 0.4$ a | $16.7 \pm 1.2$ ab | $14.7 \pm 0.9$ a |
| PDMA30 | $8.1 \pm 0.6$ a | $19.2 \pm 1.1$ a | $15.4 \pm 0.8$ a |
| Fe-EDDHA 30 | $6.7 \pm 0.4$ ab | $13.9 \pm 1.2$ bc | $13.2 \pm 0.8$ a |

### Effects of PDMA on seed quality parameters

The analysis of total protein and oil contents in seeds under various treatments revealed significant variations. Total oil content was not affected by calcareous-alkaline conditions (Table 3). However, treatments with 3 μM PDMA and 30 μM Fe-EDDHA led to similar significant increases in seed total oil content by 9% and 6% (p < 0.05), respectively, compared to both the Control (+) and Control (-) conditions. Additionally, 15 μM and 30 μM PDMA applications significantly increased seed total oil content by 16% and 19% (p < 0.05), respectively, compared to the Control (+) and Control (-) conditions. In contrast to the total oil content, seed total protein content significantly decreased by 14% (p < 0.05) when plants were grown under calcareous-alkaline conditions. Although both PDMA and Fe-EDDHA applications increased seed total protein content compared to Control (-) conditions, even the highest dosages were insufficient to restore protein content to Control (+) levels. Moreover, treatment with 30 μM PDMA increased seed total protein content by 7%, while 30 μM Fe-EDDHA treatment increased it by only 2% compared to Control (-) conditions. These results suggest that PDMA can be used to increase the protein and oil content of seeds in plants grown under calcareous-alkaline conditions. However, higher dosages of PDMA are necessary to fully restore seed protein content to Control (+) levels.

**Table 3.** Total protein and oil contents in the seeds. Data are represented as means ± SEM (n = 3). Different letters indicate a significant difference (p < 0.05) using ANOVA and Tukey’s Tests.

| **Treatments** | **Total oil content**<br>**(%)** | **Total protein content**<br>**(%)** |
| --- | --- | --- |
| C + | 19.60 ± 0.07 c | 40.30 ± 0.07 a |
| C - | 19.60 ± 0.20 c | 34.70 ± 0.02 d |
| PDMA3 | 21.50 ± 0.03 b | 35.30 ± 0.06 cd |
| PDMA15 | 22.80 ± 0.03 a | 36.60 ± 0.11 b |
| PDMA30 | 23.30 ± 0.11 a | 37.20 ± 0.04 b |
| Fe-EDDHA 30 | 20.70 ± 0.11 b | 35.40 ± 0.13 c |

Micronutrient concentrations in the seeds from each treatment group were also examined (Table 4). Interestingly, high soil pH did not significantly affect seed Fe concentrations, although the application of 30 μM Fe-EDDHA reduced seed Fe content by 14% (p < 0.05). In contrast, increasing PDMA dosages resulted in significantly higher Fe concentrations in seeds compared to Control (+). Among the micronutrients, only seed Mn concentrations were significantly lower under Control (-) conditions compared to Control (+), while Zn and Mg concentrations remained similar. However, the application of increasing PDMA dosages significantly elevated Zn, Mn, and Mg concentrations in seeds relative to Control (-) conditions. Specifically, 30 μM PDMA treatment increased seed Fe, Zn, Mn, and Mg concentrations by 39.8%, 62.3%, 66.6%, and 56.7%, respectively, compared to Control (-). In comparison, the 30 μM Fe-EDDHA treatment increased Zn, Mn, and Mg concentrations by only 28.8%, 39.7%, and 28.7%, respectively, relative to Control (-). These findings demonstrate that PDMA application not only enhances plant growth and yield but also improves seed quality by increasing micronutrient concentrations. This suggests that PDMA can serve as an effective alternative to conventional Fe fertilizers, offering a promising method for agronomic biofortification.

**Table 4.** Fe, Zn, Mn, and Mg concentrations in the seeds. Data are represented as means ± SEM (n = 6). Different letters indicate a significant difference (p < 0.05) using ANOVA and Tukey’s Tests.

| Treatments | Fe ($\mu\text{g/g DW}$) | Zn ($\mu\text{g/g DW}$) | Mn ($\mu\text{g/g DW}$) | Mg ($\mu\text{g/g DW}$) |
| --- | --- | --- | --- | --- |
| C + | $14.9 \pm 0.3$ b | $17.3 \pm 0.3$ c | $11.1 \pm 0.2$ c | $1000 \pm 31.6$ bc |
| C - | $13.8 \pm 0.3$ bc | $19.4 \pm 0.7$ c | $9.3 \pm 0.4$ d | $893 \pm 22.8$ c |
| PDMA 3 | $14.2 \pm 0.5$ bc | $18.0 \pm 0.2$ c | $10.0 \pm 0.2$ cd | $870 \pm 30.9$ c |
| PDMA 15 | $18.6 \pm 0.2$ a | $25.5 \pm 0.3$ b | $14.9 \pm 0.2$ a | $1350 \pm 37.8$ a |
| PDMA 30 | $19.3 \pm 0.5$ a | $31.5 \pm 0.8$ a | $15.5 \pm 0.4$ a | $1400 \pm 31.6$ a |
| Fe-EDDHA 30 | $12.8 \pm 0.3$ c | $25.0 \pm 0.4$ b | $13.0 \pm 0.2$ b | $1150 \pm 13.6$ b |

## Discussion

Synthetic chelates, such as EDTA, DTPA, and EDDHA, are often used to increase the solubility of divalent metal cations in the soil such to make them more available for plant uptake. Therefore, they are often used to alleviate the negative symptoms of IDC in plants. Fe-EDDHA is commonly considered the most stable of these chelates in calcareous soils due to its stronger affinity for Fe^3+^. The studies in soybean showed that soil application of these chelates ameliorates IDC and enhance the yields when grown under calcareous soils, when particularly applied before the onset of the chlorosis. However, application of high dosages of these chelates, especially Fe-EDDHA, is required to alleviate the IDC symptoms and remove the yield losses under calcareous soils because Fe fertilizers tend to become immobilized in the soil after application Due to the alkaline pH of calcareous soils (Abadía et al., 2011). To address these issues, novel adjuvants are needed to increase the Fe fertilizer efficiency.

Recently, a promising Fe chelator named PDMA was developed through the modification of DMA, a natural phytosiderophores (Suzuki et al., 2021). PDMA boasts low production costs and enhanced stability, demonstrating its efficacy in enhancing Fe absorption in rice, cucumber, and peanut plants grown hydroponically or on calcareous substrates (Suzuki et al., 2021; Ueno et al., 2021; Wang et al., 2023). Our research extends this understanding by showing that PDMA positively impacts Fe nutrition, increases soybean yield, and enhances seed quality parameters in calcareous soil under greenhouse conditions.

Our findings reveal that the application of PDMA significantly mitigates Fe deficiency symptoms in soybean plants (Figures 1 and 2). As Fe is crucial for chlorophyll synthesis (Kobayashi et al., 2019), we observed a notable increase in the chlorophyll index and total chlorophyll content with increasing PDMA dosages under alkaline conditions (Figure 1). This trend aligned with an increase in the leaf area, demonstrating a dose-dependent effect of PDMA application on leaf expansion and alleviation of Fe-deficiency chlorosis. Consequently, soybean root growth was enhanced following PDMA treatment (Figure 2), consistent with previous observations in rice (Suzuki et al., 2021), cucumber (Ueno et al., 2021), peanut (Wang et al., 2023) and maize (Suzuki et al., 2024). We demonstrated an interesting response of IDC-sensitive soybean to Fe fertilizers in the roots and shoots. While the application of PDMA increased the root lengths, these low dosages did not alter the shoot lengths (Figure 2). In contrast, Fe-EDDHA application did not significantly enhance both the shoot and root lengths as much as PDMA did when applied in the same dosage. This observation was overestimated by the previous studies (Suzuki et al., 2021; Ueno et al., 2021; Wang et al., 2023), suggesting that PDMA treatment at low dosages can affect the roots more than the shoots.

Despite the use of lower PDMA amounts in our study compared to the previous studies (Suzuki et al., 2021; Ueno et al., 2021; Wang et al., 2023), we observed similar trends in plant development, suggesting that PDMA remains effective even at lower application rates in dicots. Notably, our research highlights PDMA’s ability to enhance yield and micronutrient concentrations in soybean seeds, roots, and leaves (Tables 1-2, 4 and Figure 5), indicating its significant potential to improve both seed yield and quality under alkaline conditions. It is important to develop novel fertilizers or fertilizer adjuvants that can alleviate the IDC symptoms while increasing the yield and quality simultaneously (Maqbool et al., 2020). Although Fe-EDDHA application was shown to be as efficient as Fe–citrate in peanut (Chen et al., 2016) and soybean (Gamble et al., 2014), Fe-PDMA alleviated the IDC symptoms more effectively than Fe-EDDHA by increasing Fe concentrations in the aboveground tissues of both *Poaceae* (Suzuki et al., 2021; Suzuki et al., 2024) but also dicots, such as cucumber, pumpkin (Ueno et al., 2021), and peanut (Wang et al., 2023). Interestingly, our results prove that metal-free PDMA application is enough to alleviate the aboveground IDC symptoms in soybean in calcareous soil.

Our results show that the application of higher dosages of PDMA and Fe-EDDHA effectively mitigates the negative effects of alkalinity-based Fe deficiency on FCR activity, bringing it back to levels observed under control conditions (Figure 3a). A similar trend was also observed in the expression of known Fe uptake genes in the roots of soybean plants where PDMA application is selectively more effective in addressing Fe deficiency responses than does Fe-EDDHA at the molecular level in soybean roots (Figure 4). Interestingly, PDMA application increased the ability of plants to acidify the rhizosphere, thereby increasing the Fe bioavailability in calcareous soils (Figure 3b). Albeit our exciting results of the FCR activity, gene expressions and acidification, they do not overestimate the fact that Fe-EDDHA is a more effective Fe chelator than PDMA for enhancing root and leaf Fe concentrations, particularly in plants grown under calcareous-alkaline conditions (Figure 5). These results may emphasize the importance of application of PDMA together with Fe-EDDHA at equal but lower concentrations for a better response.

We showed that PDMA application not only increases uptake and accumulation of Fe, but also other divalent metal cations, such as Zn, Mn and Mg (Table 1). Although Fe-EDDHA could increase the Fe concentrations in the roots and trifoliate leaves significantly more than PDMA, 30 μM PDMA application caused the highest Zn and Mg concentrations in the roots and trifoliate leaves, indicating the PDMA application not only enhances Fe uptake but also some other divalent metals. These results can be explained by differential expression of *IRT1*, which can transport divalent metals such as Fe^2+^ and Zn^2+^ (Cointry and Vert, 2019). PDMA is biodegradable, is adapted to both strategy I and II Fe uptake systems, and chelates both Fe and Zn; therefore, it appears to be a promising micronutrient fertilizer.

Furthermore, our results demonstrate that PDMA application can match or surpass the seed Fe concentrations and yield values of Control (+) and even compete with Fe-EDDHA fertilizer (Tables 2, 4). This suggests that PDMA, even without combined with Fe, can positively influence plant physiological parameters and compete with Fe-containing fertilizers (Gamble et al., 2014; Gülser et al., 2019; Suzuki et al., 2021). This observation may be related to the effectiveness of PDMA in increasing the Fe bioavailability in calcareous soils by Fe-chelate solubilization without altering the pH of the potting mix (Supplementary Table 1). Indeed, similar to our results, it was shown that 40 µM PDMA application could enhance the available Fe in the soil by 20% (Wang et al., 2023).

Compared to conventional Fe fertilizers, our findings suggest that PDMA can effectively chelate insoluble Fe in the soil, promoting Fe nutrition without the need for additional Fe fertilization, thus promoting sustainable soil usage (Colombo et al., 2014). Such metal-free fertilizers efficiently utilize residual nutrients in natural soil, contributing positively to the principles of sustainable agriculture (Bouis and Welch, 2010).

In field applications, synthetic Fe chelates like Fe-EDDHA are commonly used to enhance plant Fe nutrition. However, their high cost often limits their use to cash crops (Briat et al., 2015). Moreover, synthetic chelates can persist in the soil, posing environmental concerns. As a result, there’s been a shift towards organic Fe fertilizers derived from humic acid and amino acids, which are more environmentally friendly but may lack stability, particularly in alkaline soils (Saini et al., 2016; Clemens, 2019). In contrast, PDMA offers a stable and environmentally safe alternative to synthetic chelates, avoiding pollution issues associated with them (Suzuki et al., 2021). This fact does not neglect any potential toxicity levels associated with PDMA accumulation by plants, which should be studied further. However, its application at very low dosages (such as 3 or 15 µM) seems that it does not cause any toxicity issues in soybean plants and still partially alleviates IDC symptoms under calcareous-alkaline conditions.

Another noteworthy finding from our study is that while traditional Fe-chelate-based Fe fertilizers can increase yield values in dicots, the recommended dosages by commercial companies are often excessive. Our research indicates that comparable yield values can be achieved with significantly lower amounts of PDMA. In fact, PDMA and similar compounds can potentially replace conventional Fe fertilizers without adding additional Fe to the soil, thus minimizing environmental damage. In a previous study, application of FeSO_4_ at an Fe/soil ratio of 15:1000 caused a significant increase in soil pH and total phosphate levels while significantly decreasing the total nitrogen, ammonium and nitrate levels (Fazl et al., 2024). These changes in soil nutrient levels, particularly a sharp increase in the sulfur content, in turn, altered the bacterial and fungal community structure of the soil. Although this study did not investigate the effect of PDMA application on microbial diversity and interactions in the soil, it is noteworthy to analyze the soil microbiome after PDMA application.

Regarding the relationship between nitrogen and Fe metabolisms, Fe deficiency typically reduces nitrogen fixation and amino acid production in plants. This is because Fe is crucial for the synthesis of chlorophyll and various enzymes involved in nitrogen metabolism. As such, Fe-deficient plants often show reduced protein levels due to impaired nitrogen assimilation (López-Millán et al., 2013; López-Pérez et al., 2024). However, PDMA treatment appears to counteract these adverse effects of Fe deficiency by improving Fe availability necessary for nitrogen metabolism. Total protein content increases with increased PDMA concentration. 30 µM PDMA application shows the second highest results after Control (+), indicating its role in enhancing the availability of Fe, which is crucial for nitrogen metabolism (Table 3). Furthermore, soybeans grown in calcareous-alkaline conditions typically exhibit lower protein and oil content due to Fe absorption difficulties, with these nutrient levels decreasing as soil pH increases (Chen et al., 2016; Chen et al., 2018). Yet, PDMA treatment seems to mitigate and reverse this decline, suggesting it could help maintain or enhance the nutrient balance in soybean seeds under such challenging conditions.

While PDMA showed superior performance to Fe-EDDHA in certain growth and nutrient accumulation metrics, its specific effects on different micronutrients suggest a unique mechanism of action that warrants further investigation. The significant reduction in FCR activity and the modulation of Fe uptake gene expression also highlight PDMA’s potential in fine-tuning iron homeostasis. Future research should focus on field trials to validate the effectiveness of PDMA under more variable environmental conditions. Additionally, exploring the long-term impacts of PDMA on soil health and its interaction with other plant nutrients could provide insights into optimizing its use in sustainable agriculture. Exploring the molecular mechanisms underlying PDMA’s effects could further refine its application and expand its use across different crop systems.

## Conclusion

This study demonstrates that PDMA effectively alleviates Fe deficiency symptoms in a soybean cultivar susceptible to Fe deficiency, grown under calcareous-alkaline conditions. PDMA application significantly improved Fe uptake, translocation, and storage within the plant, leading to increased chlorophyll content, root and shoot growth, and enhanced overall plant development. Importantly, PDMA not only boosted Fe nutrition but also promoted the accumulation of essential micronutrients such as Zn, Mn, and Mg, particularly at higher concentrations (30 μM). These enhancements resulted in improved soybean yield parameters, including pod and seed number, as well as 100-seed weight. Furthermore, seed quality was elevated, with higher protein and oil content observed following PDMA treatment. These findings highlight PDMA’s significant potential as a highly effective Fe supplement and growth promoter for agricultural practices, particularly in Fe-deficient conditions.

## Acknowledgments

This research is supported by the grant provided by AICHI STEEL CORPORATION. The authors thank Assoc. Prof. Seçkin Eroğlu for his critic evaluation of the manuscript before submission.

## Author Contribution Statement

ZK and EA conceived and designed research. MS synthesized the PDMA. ZK and AM conducted the experiments. ZK, AM and EA analyzed the data. ZK and EA wrote the manuscript. EA and MS edited the manuscript. All authors read and approved the manuscript.

