## Supplementary material for "Proline-2’-deoxymugineic acid (PDMA) increases seed quality and yield by alleviating iron deficiency symptoms in soybean under calcareous-alkaline conditions": Suppl_table_1

**Supplemental Table 1.** Properties of the soil used in the study. Samples were collected from 0-30 cm. Data represents average ± SEM (n=3).

| **Soil parameters** | **Before the experiment** | **After the experiment** |
| --- | --- | --- |
| Sand (%) | 11.25 ± 1.3 | 13.7 ± 1.4 |
| Silt (%) | 20.50 ± 1.7 | 20.75 ± 1.5 |
| Clay (%) | 68.25 ± 3.5 | 66.55 ± 2.4 |
| pH | 8.32 ± 0.11 | 8.24 ± 0.14 |
| EC (mS/cm) | 0.26 ± 0.04 | 0.25 ± 0.02 |
| CaCO_3_ (%) | 22.5 ± 0.98 | 21.45 ± 1.03 |
| Organic matter (%) | 1.92 ± 0.12 | 1.95 ± 0.14 |
| Total N (%) | 0.12 ± 0.02 | 0.14 ± 0.03 |
| Available P (mg/kg) | 22.50 ± 1.4 | 21.45 ± 1.3 |
| Available Fe (mg/kg) | 2.65 ± 0.56 | 3.12 ± 0.52 |
| Available Zn (mg/kg) | 0.95 ± 0.08 | 1.10 ± 0.04 |
| Available Mg (mg/kg) | 5150.12 ± 105.14 | 5225.14 ± 114.45 |
| Available Mn (mg/kg) | 5.45 ± 0.89 | 6.17 ± 0.84 |
| Available Cu (mg/kg) | 0.11 ± 0.04 | 0.12 ± 0.02 |
| Extractable Fe (mg/kg) | 1.89 ± 0.11 | 2.85 ± 0.09 * |

* represents statistical difference between the groups according to Student’s t-test (p < 0.05).
